# Skin development and metamorphic re-arrangement in *Arthroleptella villiersi* Hewitt, 1935 reveals distinct heterochronic patterns in anurans with different life histories

**DOI:** 10.1101/2025.04.06.647201

**Authors:** Benjamin Naumann, Susan Schweiger, Hendrik Müller

## Abstract

**Background:** The evolvability of ontogenetic trajectories, driven by modular developmental processes and their evolutionary repatterning, plays a crucial role in the diversity of animal diversity. Anurans offer a model for exploring how developmental repatterning, including the loss of larval features and heterochronic shifts of metamorphic processes lead to the diversity of, often terrestrialized, life histories. Ancestrally, anurans exhibit an aquatic larva (tadpoles) that differs tremendously from their semi-terrestrial adult forms. The transition from this tadpole to the adult is a thyroid hormone-dependent metamorphosis that results in the rearrangement of almost all organ systems. One of the organ systems that experiences the most extensive rearrangements is the skin. Is the ancestral pattern of metamorphic skin rearrangement altered in anurans with a completely terrestrial tadpole phase?

**Results:** Skin development and metamorphic rearrangement in the terrestrial tadpole of *Arthroleptella villiersi* is investigated based on histological sections. It follows the major pattern observed in anurans with an aquatic tadpole phase and correlates with thyroid gland maturation. Neuromasts develop and are present during a short developmental time. Both, metamorphic skin rearrangement and neuromast degeneration correlate with histological thyroid gland maturation. A few minor heterochronic shifts are detectable compared to aquatic indirect and terrestrial direct developing anurans.

**Conclusion:** Skin development and metamorphic rearrangement in anurans with different reproductive modes is very similar and correlates with thyroid gland maturation indicating that skin development in anurans might be constrained by thyroid hormone activity. Neuromasts are the major exception being present in aquatic and terrestrial indirect developing species but not in terrestrial direct developing anurans. Heterochronic shifts detected during skin development indicate that at least the differentiation of multicellular gland complexes might represent a developmental module similar to neuromasts.

## 1. Introduction

The evolvability of ontogenetic trajectories, their capacity to produce heritable variants, is key to the evolution of the diversity of animal life histories (Gerhart and Kirschner, 1997; Von Dassow and Munro, 1999). Evolvability is enhanced by the modular organization of developmental processes, enabling alterations within discrete developmental entities (modules) independent from other such modules and thereby maintaining overall organismal integrity. Evolutionary alterations can be described as developmental repatterning and are often categorized as heterotypy (change in type), heterotopy (change in space), heterometry (change in amount) and heterochrony (change in time) (Arthur, 2021). The question arising from this framework is whether certain developmental repatterning processes are more prevalent in changes of ontogenetic trajectories and have larger explanatory value for the evolution of the diversity of animal life histories.

Extant amphibians - caecilians (Gymnophiona), salamanders (Caudata) and frogs (Anura) – are a well-suited animal group to approach this question. Their ancestral life history is characterized by an indirect reproductive mode, including an aquatic larva that transforms via a thyroid hormone-dependent metamorphosis into a semi-terrestrial adult (Duellman and Trueb, 1994). Starting from this, all three amphibian groups have evolved diverse derived life histories, including aquatic, semi-terrestrial, and fully terrestrial reproductive modes, with varying degrees of larval reduction up to its complete loss in direct-developing taxa (Liedtke et al., 2022). With more than 7.800 described species (amphibiaweb.org, 15.03.2025), anurans comprise the evolutionarily most successful amphibian group in terms of species richness. This evolutionary success is largely attributed to their diverse indirect life histories including highly specialized free-swimming and feeding larvae known as tadpoles (Altig and McDiarmid, 1999). Within anurans, terrestrial reproductive modes, including the reduction (terrestrial-indirect development) or even complete loss of the free-living tadpole phase (direct development), have evolved independently more than 20 times (Figure 1(A)) (Liedtke et al., 2022). Among these terrestrial reproductive modes, the evolution of direct development has received most attention. Previous studies revealed evolutionary losses of tadpole-specific features such as the larval head skeleton, associated musculature, adhesive glands, lateral line organs, and coiled intestine (Hanken et al., 1997; Naumann et al., 2020; Townsend and Stewart, 1985). Other tadpole-specific features such as a long tail, an epidermal covering of gills and forelimbs (operculum) and a two-layered epidermis with unicellular glands still develop (Callery, 2000; Callery and Elinson, 2000; Naumann et al., 2020; Townsend and Stewart, 1985). These features experience a metamorphic re-arrangement early in development and, together with features that appear at an already mid-metamorphic state, have led to the hypothesis that large aspects of the different ontogenetic trajectories underlying anuran life history diversity evolved via heterochrony (Gould, 1977). Pre-displacement of pituitary and thyroid gland maturation, the so-called brain-pituitary-thyroid-axis, has been proposed as the main event causing the heterochronic shifts of many metamorphic and adult features (Jennings and Hanken, 1998; Laslo et al., 2019; Morgan et al., 1989; Naumann et al., 2020). However, testing the hypothesis on heterochrony as the main pattern underlying the diversity of anuran life histories and reproductive modes requires detailed comparative data from anurans with diverse life histories and reproductive modes. These data, including the developmental timing of various organ systems, can then be compared using formalized techniques like heterochrony plots to reliably identify patterns of temporal displacement and co-dissociation (Schlosser, 2001).

**Figure 1.**
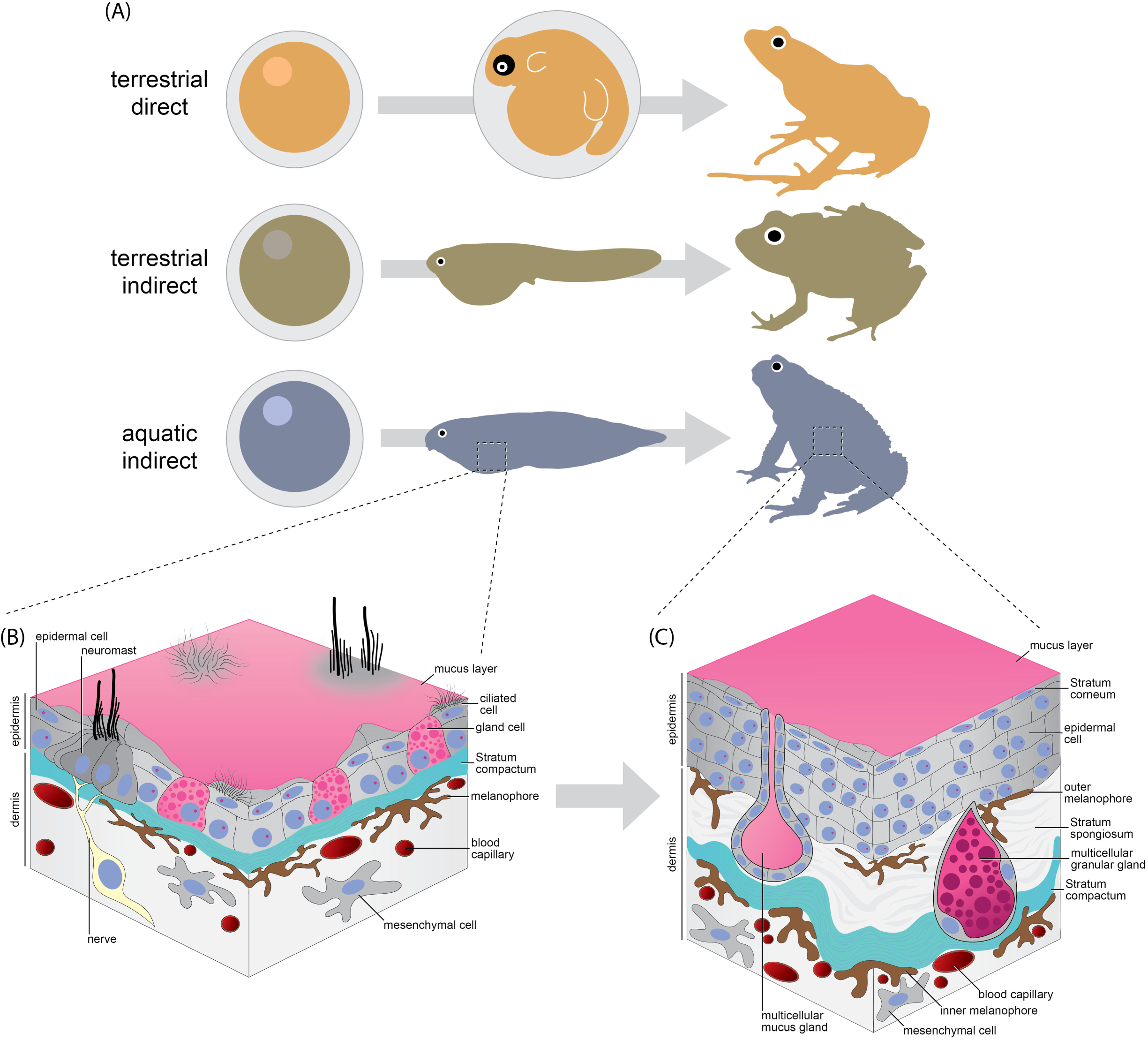
(A) Schematic illustrations of different reproductive modes in Anura. For each reproductive mode, an egg is depicted on the left, an embryo/tadpole in the middle and an adult on the right. (B) Schematic illustration of the skin of a free-living, aquatic tadpole. (C) Schematic illustration of the skin of a postmetamorphic anuran.

To contribute to the aforementioned program, we investigated the timing of skin and thyroid gland development in the terrestrial indirect-developing frog *Arthroleptella villiersi* Hewitt, 1935 (Pyxicephalidae). In *A. villiersi,* eggs are laid terrestrially beneath vegetation, in loose soil, or organic matter, and are surrounded by a moist jelly mass. Non-feeding (endotrophic) tadpoles hatch from these eggs at an early “metamorphic” stage, and continue their development within the jelly nest until metamorphosis is completed and small froglets emerge from the nest (de Villiers, 1929). Thus, the life history of this species combines features of both terrestrial direct-development and aquatic-indirect development including an apparent metamorphosis (de Villiers, 1929; Morgan et al., 1989). While many organs are rearranged during anuran metamorphosis, the skin exhibits one of the most extensive rearrangements and has been suggested as a model system to investigate the evolutionary alteration of ontogeny in anurans (Naumann et al., 2020; Yoshizato, 1992). The skin of pre-metamorphic tadpoles is composed of a two-layered epidermis that includes unicellular glands and an outer mucus layer.

Neuromasts, organized in distinct lateral lines, are embedded within this larval epidermis. The underlying dermis is a mesenchyme that includes a stratum compactum and melanophores (Figure 1 (B)). Entering pro-metamorphosis, most epidermal cells degenerate, except for those connected to the basal lamina, and begin to proliferate again during the metamorphic climax. In most species, neuromasts degenerate together with the larval epidermis and only persist during the post-metamorphic phase in taxa with aquatic adults such as *Xenopus* (Mohr and Görner, 1996). The post-metamorphic epidermis forms a five- to seven-layered epithelium with an outer keratinized stratum corneum and various types of multicellular glands. The dermis contains a stratum spongiosum and both an outer (located distal to the stratum compactum) and an inner melanophore layer (positioned proximal to the stratum compactum) (Figure 1 (C)) (Fox, 1986; Fox and Whitear, 1990; Naumann et al., 2020). Morgan and colleagues (1989) described a conserved overall pattern of skin development and metamorphic rearrangement in *Arthroleptella lightfooti* (Boulenger, 1910) compared to *Xenopus laevis* Daudin, 1802, but did not report the exact timing of specific developmental events, such as the induction, maturation and degeneration of neuromasts or the differentiation of different multicellular gland complexes. However, these data are necessary for a detailed heterochrony analysis. The aim of this study is to investigate skin development and metamorphic repatterning in *A. villiersi* in detail and analyse it using a heterochrony plot against the aquatic indirect-developing bufonid *Rhinella arenarum* (Hensel, 1867) assumed to represent a more ancestral pattern. We also analyse skin development in the terrestrial direct-developing *Arthroleptis wahlbergii* Smith, 1849 in comparison with *R. arenarum* and *A. villiersi*, to evaluate heterochronic patterns of skin development in anurans with different life histories.

## 2. Results

### 2.1 Development of the pigmentation pattern

**Stage 6 (Figure 2(A)):** Only a very few scattered melanophores are detectable in the skin of the dorsal head and trunk at this developmental stage.

**Figure 2.**
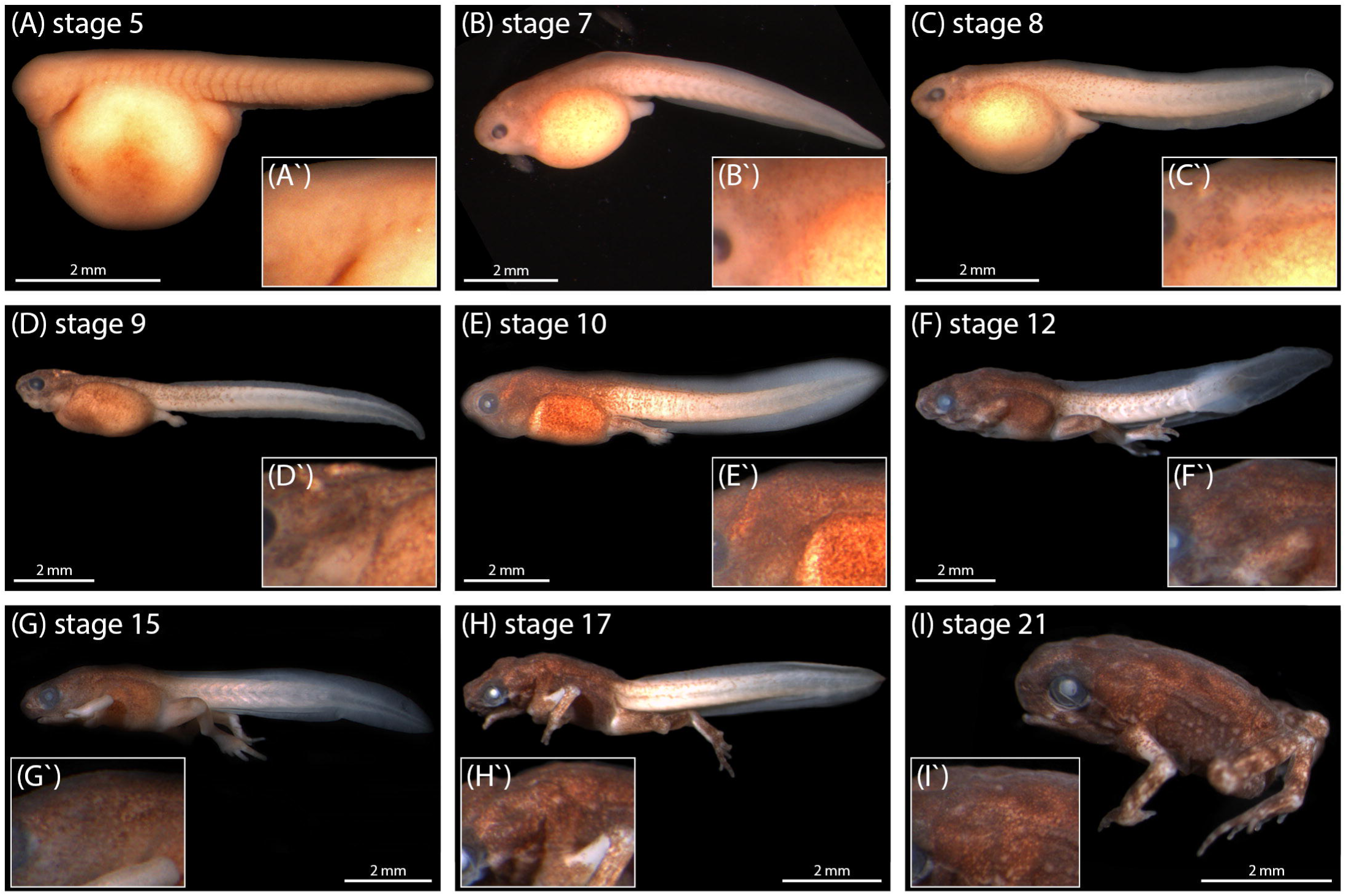
Photographs of different developmental stages of *Arthroleptella villiersi* in lateral position to illustrate the developing pigmentation pattern. (À-Ì) show a close up of the posterior head region of each main photograph.

**Stage 7 (Figure 2(B)):** Melanophores are present all over the skin of the head and trunk. Most of these melanophores exhibit a stellate morphology with a few short extensions and a broad and flattened central region. Their density is highest at the dorsal midline and decreases ventrolaterad until the level of the eye on the head and the dorsal end of the pronephric system in the trunk. Ventral to that level, melanophores are concentrating in the skin lateroventrally to the eye and overlaying the pronephric system. From that level on, melanophore density decreases again ventrolaterad. No melanophores can be seen in the tail skin.

**Stage 8 (Figure 2(C)):** In the trunk, melanophores are present down to a more ventral level compared to the previous developmental stage. A very few of them reach the ventral midline of the trunk. The anterior half of the tail also exhibits melanophores. Their concentration is highest dorsally, decreasing ventrolaterad.

**Stage 9 (Figure 2(D))**: Melanophore density has increased tremendously compared to the previous developmental stage. A higher number of them reaching the ventral midline of the trunk. Pigmentation of the tail has extended to the anterior two thirds to three quarters of its length.

**Stages 10 to 14 (Figures 2(E - F))**: Density of melanophores increases during this developmental period. In all these stages pigmentation of the tail skin only reaches up the anterior three quarters of the tail length. The posterior quarter of the tail shows only a very few separated melanophores and appears almost unpigmented.

**Stages 15 to 22 (Figures 2(G – I))**: Density of melanophores increases further, reaching a pattern comparable to stage 20-juveniles at around stage 15. Pigmentation of the tail fades in parallel with its degeneration.

### 2.2 Development of skin histology

**Stage 5 to 6 (Figure 3(A)): Head** – The epidermis is single-layered with a sometimes-indistinct border between the underlying cells. Epidermal cells are large and cuboid with spherical nuclei, large vacuoles and a high number of intracellular yolk droplets (Figure 3(A1)). **Trunk** – Dorsally, the epidermis is mostly single-layered with a few patch-like two-layered areas. Epidermal cells of head and trunk are similar (Figure 3(A2-3)). Ventrally, the epidermis is single-layered. The cells are more flattened compared to the dorsal trunk region. They exhibit large spherical nuclei, large vacuoles, intracellular yolk droplets and are clearly separated from the underlying yolk sac (Figure 3(A3)). **Tail** – The epidermis is single-layered and cells are similar compared to the trunk (Figure 3(A4)).

**Figure 3.**
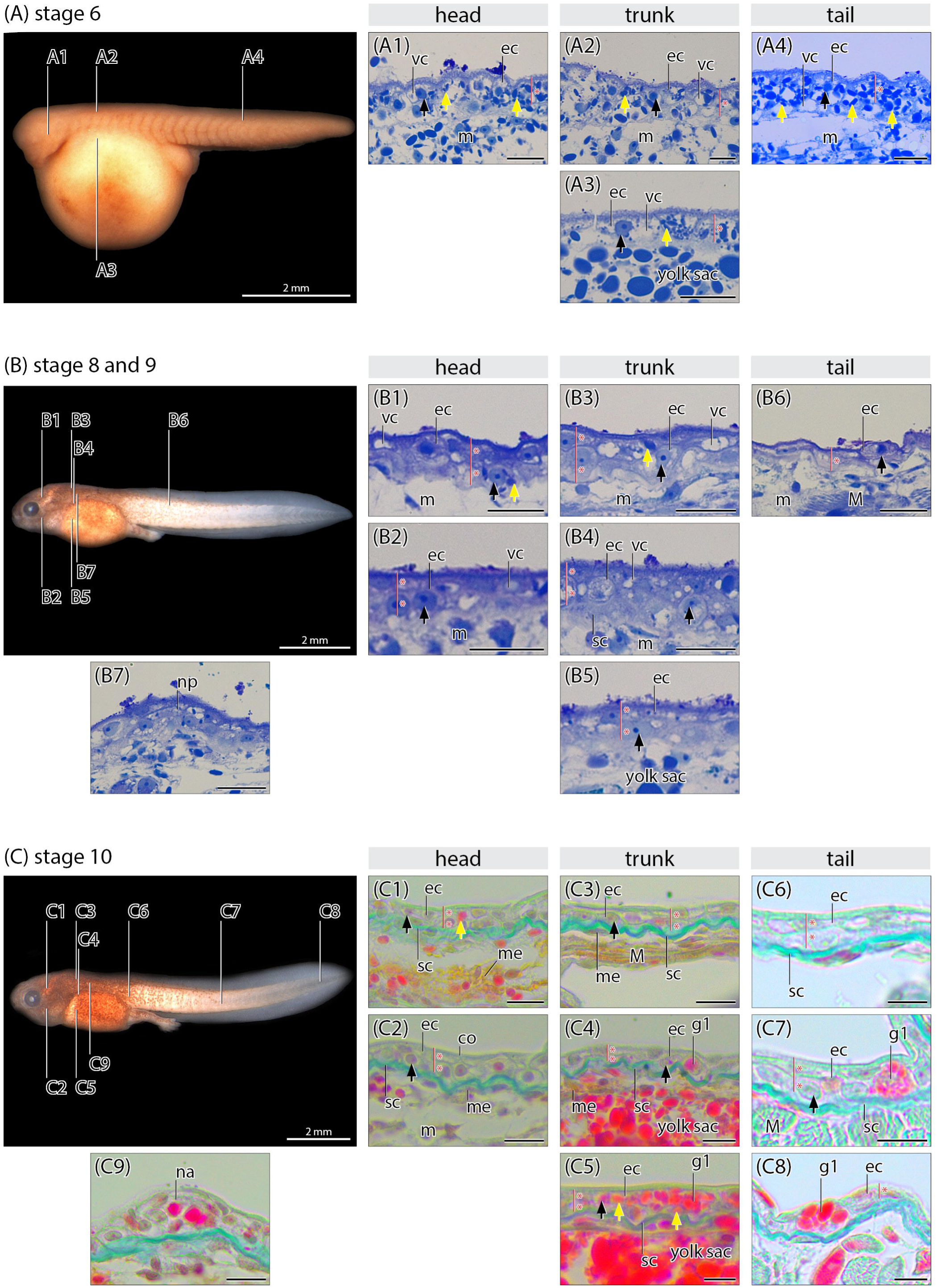
Histological cross sections of the skin of *Arthroleptella villiersi* at different developmental stages. Yellow arrows indicate intracellular yolk droplets. Black arrows indicate epidermal nuclei. The number of epidermal cellular layers is indicated by a red line and asterisks. Scale bars in histological sections equal approximately 200 μm. ec, epithel cell; g, multicellular gland progenitor; g1, unicellular gland; M, muscle; m, mesenchyme; mc, mantle cell; me, melanophore; np, neuromast placode; sc, Stratum compactum; sp, Stratum spongiosum; su, support cell; vc, vacuole.

**Stage 8 and 9 (Figure 3(B)): Head** – Dorsally, the epidermis is two-layered. Ventrally, most of the areas are two-layered except for a few single-layered patches. The epidermis is separated from the underlying cells by a sharp boarder. However, a stratum compactum cannot be detected in this developmental phase. The cells are still large and cuboid with spherical nuclei. The number of vacuoles and intracellular yolk droplets is greatly reduced compared to the previous developmental stage (Figures 3(B1)). **Trunk** – The epidermis is two-layered. Dorsally, the external cell-layer is more flattened compared to the internal layer of cuboid cells (Figure 3(B2)). Medially, the two layers are more or less similar. Ventrally, both cell layers are more flattened, and nuclei show an ovoid shape. The number of vacuoles and intracellular yolk droplets is also reduced. **Tail** – The epidermis is single-layered. The cells are flattened with ovoid-shaped nuclei and almost no vacuoles and intracellular yolk droplets (Figure 3(B3)). A few patches with more densely clustered cells, slightly protruding from the rest of the epidermis, are present in the dorsal skin of the trunk and are interpreted as early neuromast placodes (Figure 3(B4)).

**Stage 10 (Figure 3(C)): Head** – Dorsally, the epidermis is two-layered and a dense area of melanophores underlies a thin stratum compactum (internal melanophores). Ventrally, a few patch-like single-layered areas are still present overlying a thin stratum compactum. The number of melanophores decreases towards the ventral midline. No vacuoles and only very few scattered intracellular yolk droplets are present (Figures 3(C1-C2)). **Trunk** – The epidermis is mostly two-layered. Epidermal cells are more flattened compared to previous developmental stages. Most nuclei exhibit an ovoid-shape. The stratum compactum is distinct. Neither vacuoles nor intracellular yolk droplets can be found dorsally. Laterally, their number increases, resulting in a high number of intracellular yolk droplet in the ventral epidermis. Melanophores underlying the stratum compactum are only present dorsally (Figures 3(C3-C4)). **Tail** – The epidermis is single- to two-layered. Cells in the external layer are flattened with ovoid nuclei, while cells in the internal layer are more cuboid with round nuclei. At the tail tip, cells become cuboidal. A stratum compactum is present. A few scattered gland cells are detectable in the middle area and close to the tail tip. No melanophores are recognizable (Figures 3(C5)). Neuromast anlagen, clearly recognizable by their rosette-like shape, are present at the positions of the placodes of the previous developmental stage (Figure 3(C6)).

**Stage 12 (Figure 4(A)): Head** – The epidermis is two-layered. The stratum compactum has increased in thickness. Anlagen of large multicellular glands, recognizable as large round cell clusters often sunken into the stratum compactum, are present in the dorsal area. Epidermal organization as a stratum spongiosum is first detected around multicellular gland complexes (Figures 3(D1-D2)). **Trunk** – Dorsally, the epidermis is single- to two-layered. Laterally, the epidermis is single-layered with a decreasing density of melanophores. Laterally, most areas of the epidermis are single-layered. Ventrally, the epidermis is single-layered. A dense layer of melanophores underlies the stratum compactum and a stratum spongiosum is first recognizable (Figures 3(D3-D4)). **Tail** – Cells in both epidermal layers appear more flattened compared to previous developmental stages. Anlagen of multicellular glands can be recognized. The epidermis at the posterior-most tail area and the tip is single layered. (Figures 3(D5-7)). Mature neuromasts are present in the positions of the anlagen described at the previous developmental stage. Ciliated median cells are surrounded in a rosette-like pattern by non-ciliated support cells and outer-most mantle cells. The mantle cells protrude centrally, forming an externally opening cavity that encloses the support and ciliated cells (Figure 3(D8)).

**Figure 4.**
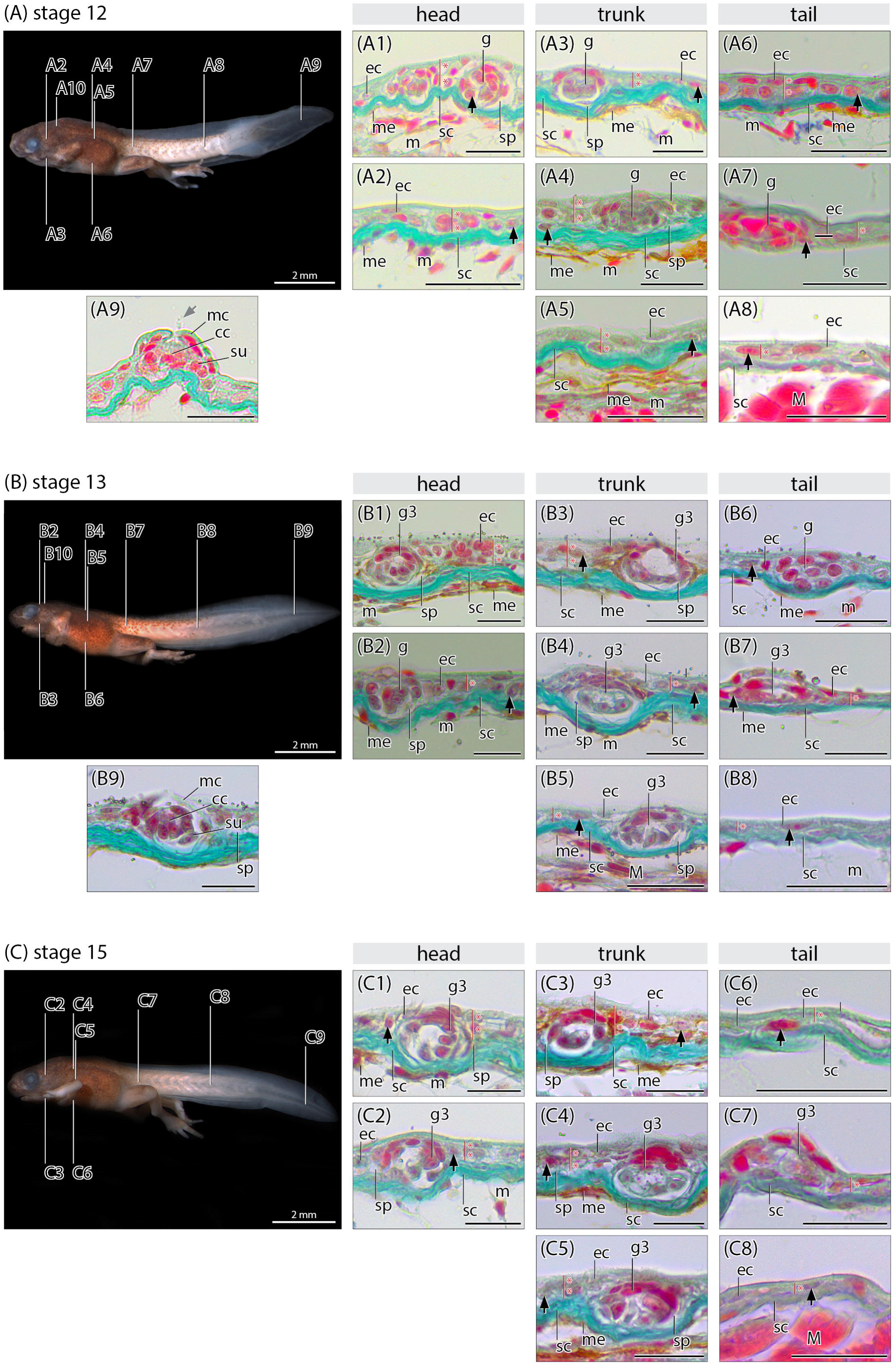
Histological cross sections of the skin of *Arthroleptella villiersi* at different developmental stages. Black arrows indicate epidermal nuclei. Grey arrows indicate the neuromast cilium. The number of epidermal cellular layers is indicated by a red line and asterisks. Scale bars in histological sections equal approximately 200 μm. cc, cilium cell; co, Stratum corneum; ec, epithel cell; g, multicellular gland progenitor; g1, unicellular gland; g2, multicellular granular gland; g3, multicellular mucus gland; M, muscle; m, mesenchyme; mc, mantle cell; me, melanophore; sc, Stratum compactum; sp, Stratum spongiosum; su, support cell.

**Stage 13 (Figure 4(B)): Head** – The epidermis is single- to two-layered with more areas being single-layered compared to the previous developmental stage. In the two-layered areas, cells of the internal cell layer have a more cuboid shape and nuclei appear more rounded indicating an increased mitotic activity. The number of multicellular glands has increased and some of them can be identified as mucus glands showing a cellular wall surrounding a large central lumen (Figure 4(A1-A2)). **Trunk** – The epidermis is single- to two-layered. In the two-layered regions, cells of the internal layer appear cuboid in shape, with round nuclei similar to the head region. Cells of the external layer are more flattened, and their nuclei are more ovoid in shape. The number of multicellular glands has increased in the dorsal and lateral skin and their differentiation appears more pronounced compared to the head (Figures 4(A3-A4)). **Tail** – The epidermis is single- to two-layered consisting of flat cells. The number of developing multicellular glands has increased. However, they appear much less differentiated compared to the multicellular glands present in the epidermis of the trunk and head (Figures 4(A5-A7)). Neuromasts are still present in regions similar to the previous developmental stage. However, no cilia are found in most of them, indicating degenerative processes comparable to the epidermis cells of the tail tip (Figure 4(A8)).

**Stage 15 (Figure 4(C)): Head** – The number of cell layers and the shape of epidermal cells is similar compared to the previous stage. The stratum compactum appears thicker, and the number of multicellular glands has increased compared to the previous stage. Most of them can be clearly identified as mucus glands (Figure 4(B1-B2)). **Trunk** – As for the head, the number of cell layers and cellular shapes are similar to the previous stage, but the stratum compactum appears thicker compared to previous developmental stages. The number and stage of differentiation of mucus glands has increased compared to the previous developmental stages (Figure 4(B3-B4)). **Tail** – The epidermis appears similar to the previous stage except that the number of multicellular glands has decreased (Figure 4(B5-B7)). Neuromasts are no longer found.

**Stage 17 (Figure 5(A)):** From this developmental stage on, the skin of the embryo very much resembles the skin of juvenile frogs at stage 20. **Head** – Number and size of mucus glands have increased, and first multicellular granular glands are present in the dorsal epidermis. Additionally, a dense layer of melanophores is present in the dorsal epidermis, lying externally to the stratum compactum (external melanophores) (Figure 5(C1)). **Trunk** – Dorsally, most areas of the epidermis are three-layered, thinning to a two- or single-layered epidermis towards the ventral midline. Similar to the epidermis of the head, the number and size of multicellular mucus glands have increased, the first multicellular granular glands are present in the dorsal region, and a dense layer of external melanophores has formed, decreasing in a ventral direction (Figure 4(C2-C3)). **Tail** – The epidermis appears mostly similar to the previous stage. The number of multicellular glands has further decreased (Figure 4(C4)).

**Figure 5.**
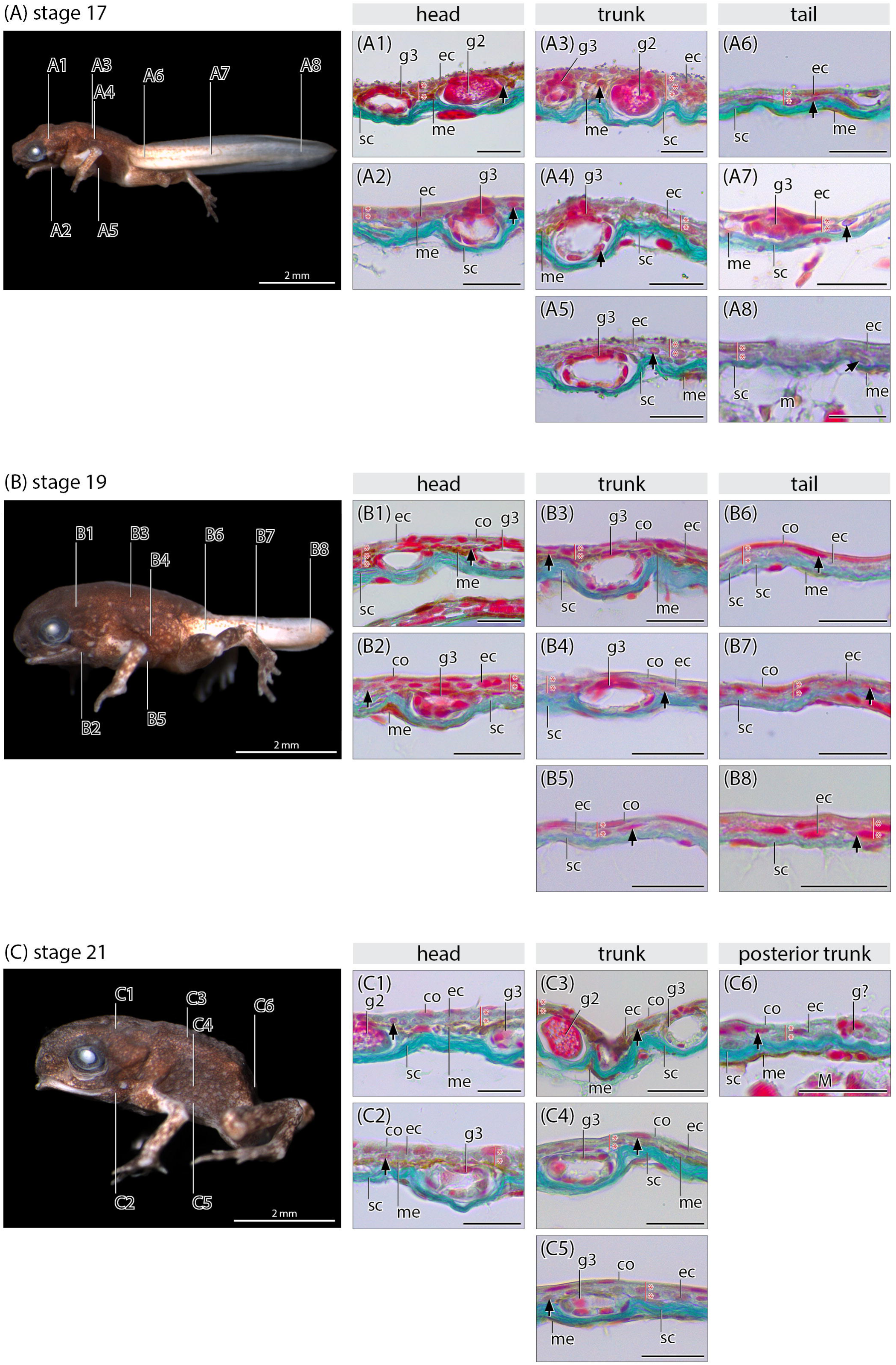
Histological cross sections of the skin of *Arthroleptella villiersi* at different developmental stages. Black arrows indicate epidermal nuclei. The number of epidermal cellular layers is indicated by a red line and asterisks. Scale bars in histological sections equal approximately 200 μm. co, Stratum corneum; ec, epithel cell; g?, degenerating gland; g2, multicellular granular gland; g3, multicellular mucus gland; M, muscle; m, mesenchyme; me, melanophore; sc, Stratum compactum; sp, Stratum spongiosum.

**Stage 19 (Figure 5(B)): Head** – In some patchy areas of the dorsal and lateral sides of the head, the dorsal epidermis is three-layered. The external cell layer is flattened and represents the stratum corneum. A slightly higher number of multicellular mucus and granular glands is present compared to previous stages (Figures 4(D1)). **Trunk** – The skin is similar compared to the previous stage except for a stratum corneum, a higher number of multicellular mucus and granular glands and an increased thickness of the stratum compactum (Figures 4(D2)). **Tail** – The skin is similar compared to the previous developmental stage (Figure 4(D3)).

**Stage 21 (Figure 5(C)): Head** – The epidermis is mostly similar to the previous developmental stage. The density of external melanophores has increased in the dorsal epidermis, while there are still none in the ventral epidermis. The number of granular glands has increased compared to previous developmental stages (Figures 4(E1)). **Trunk** – The epidermis is similar to the previous developmental stage except for a higher density of outer melanophores and an increased number of multicellular granular glands (Figures 4(E2)). **Posterior trunk** – The tail has been completely resorbed. The epidermis overlaying posterior trunk region is two-layered, exhibiting only inner melanophores (Figure 5(E3)).

### 2.3 Development of the thyroid gland

For the histological description and measurements, sections of one specimen each for developmental stage 10, 12, 13, 15, 17, 19, 21 and 22 were analysed (Figures 5 and 6).

**Figure 6.**
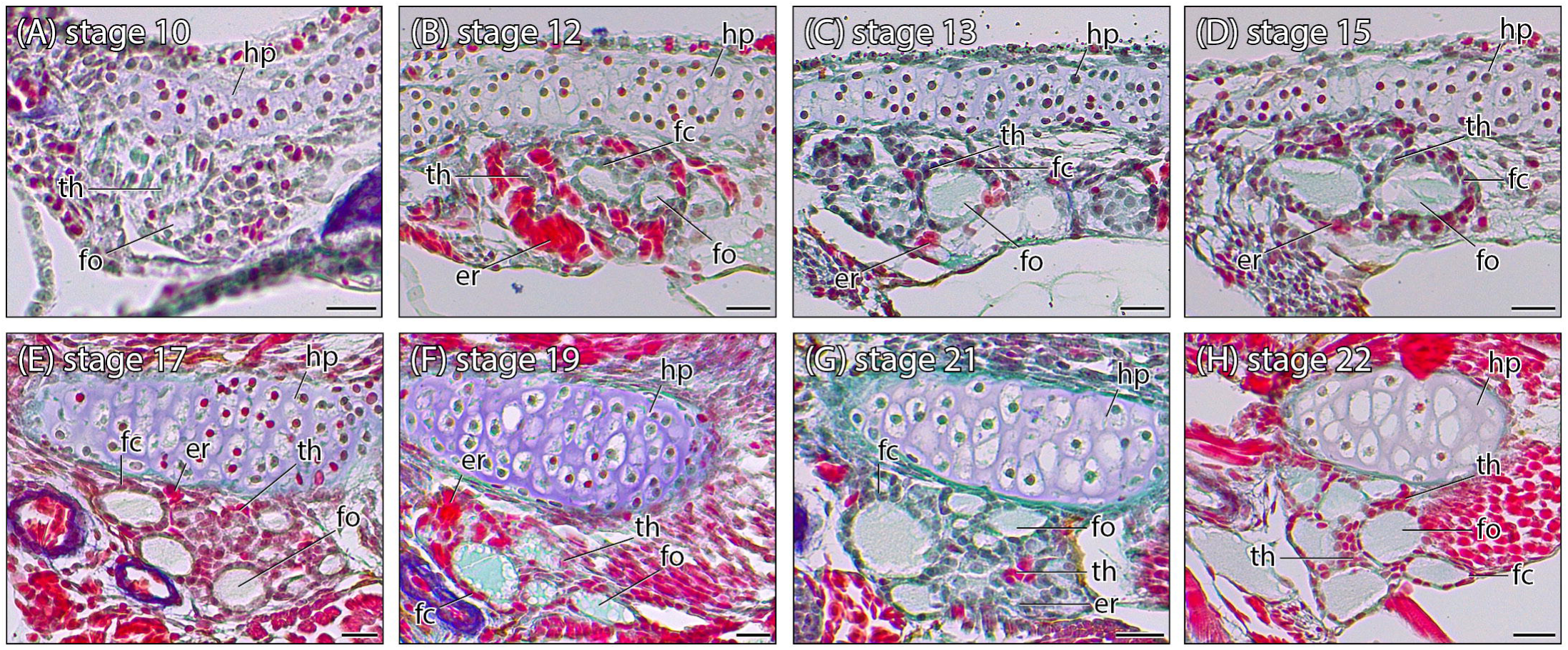
Histological cross sections of the ventral head region of *Arthroleptella villiersi* at different developmental stages showing thyroid gland development. Scale bars equal approximately 200 µm. er, erethrocytes; hp, hyobranchial plate; fc, follicle cell; fo, follicle; th, thyroid gland.

The thyroid gland of *A. villiersi* is first recognizable at stage 8 as a spherical cell condensation ventral to the right and left lateral margin of the developing hyoid plate (Figures 5(A). Follicles, showing only a loosely organized outline by follicular wall cells, are present in histological sections of a specimen at this stage. At stage 10, follicular wall cells are organized as a compact wall exhibiting cuboid cell shapes and large spherical nuclei. The presence of a high number of erythrocytes indicates an increased vascularization of the thyroid gland. Faintly stained colloid is first detectable in follicular lumina (Figures 5(B)). At stage 13, follicular wall cells are organized in a columnar pattern. Their nuclei exhibit a range of shapes from spherical to ovoid, indicating increased mitotic activity. Staining of colloid inside the follicular lumen is more intense compared to the previous developmental stage (Figures 5(C)). At stage 15, histology of the thyroid gland appears similar to the previous developmental stage (Figures 5(D)). At stage 17, follicular wall cells exhibit larger amounts of cytoplasm compared to previous developmental stages (Figures 5(E)). From this stage on, the histological organization is similar compared to stage 19, stage 21 and stage 22 and resembles the mature configuration (Figures 5(F-H)). The number of detectable erythrocytes decreases from stage 17 to stage 21, indicating a decreasing degree of vascularization and thyroid gland activity.

To estimate thyroid gland activity based on histological sections, the follicle number, single follicle volume, total follicle volume, and follicular wall cell height of the thyroid glands were measured (Figure 6; outliers are not depicted in Figures 6(A) and (C)). Values of measurements are given in Supplementary Material 1.

### 2.4 Heterochrony analysis of skin development in anurans with different reproductive modes

In a heterochrony plot, a species showing the presumably derived condition is plotted against a species representing the plesiomorphic condition (Schlosser, 2001). Therefore, data from the aquatic indirect-developing species *X. laevis* and *R. arenarum* were plotted against each other to determine the degree of similarity of the timing of developmental events and to test their use as a potential plesiomorphic species in comparison with anurans showing a derived reproductive mode (Figure 7A). The resulting heterochrony plot shows a comparable developmental timing indicated by the linear regression lines, a rough approximation of the developmental velocity of the suites, clustering closely together. However, a few single events exhibit a shift in timing summarized in Table 1. Because of the very similar timing and the higher number of available data on skin developmental events, *R. arenarum* was chosen as a species representing the plesiomorphic condition in subsequent plots. To test for heterochronic shifts of skin development connected with more terrestrial reproductive modes, data from the terrestrial indirect-developing *A. villiersi* are plotted against *R. arenarum* (Figure 7B). An additional plot was prepared for the terrestrial direct-developing *A. wahlbergii* against *R. arenarum* to test if heterochrony patterns of terrestrial endotrophic *Arthroleptella* and *Arthroleptis* are similar compared to the plesiomorphic condition (Fig. 7C). A heterochrony plot of *A. villiersi* against *A. wahlbergii* was also prepared to test the similarity of terrestrial endotrophic reproductive modes (Figure 7D). For the latter, the terms “earlier” and “later” instead of “pre-displaced” and “post-displaced” are used to regard that there is no evidence that *Arthroleptis* evolved from a terrestrial-indirect developing ancestor. A summary of the heterochronic shifts detected in the plots is given in Table 1. The timing of developmental events defining the ontogenetic phases and events within each suite and are given in Table 2 and 3.

**Figure 7.**
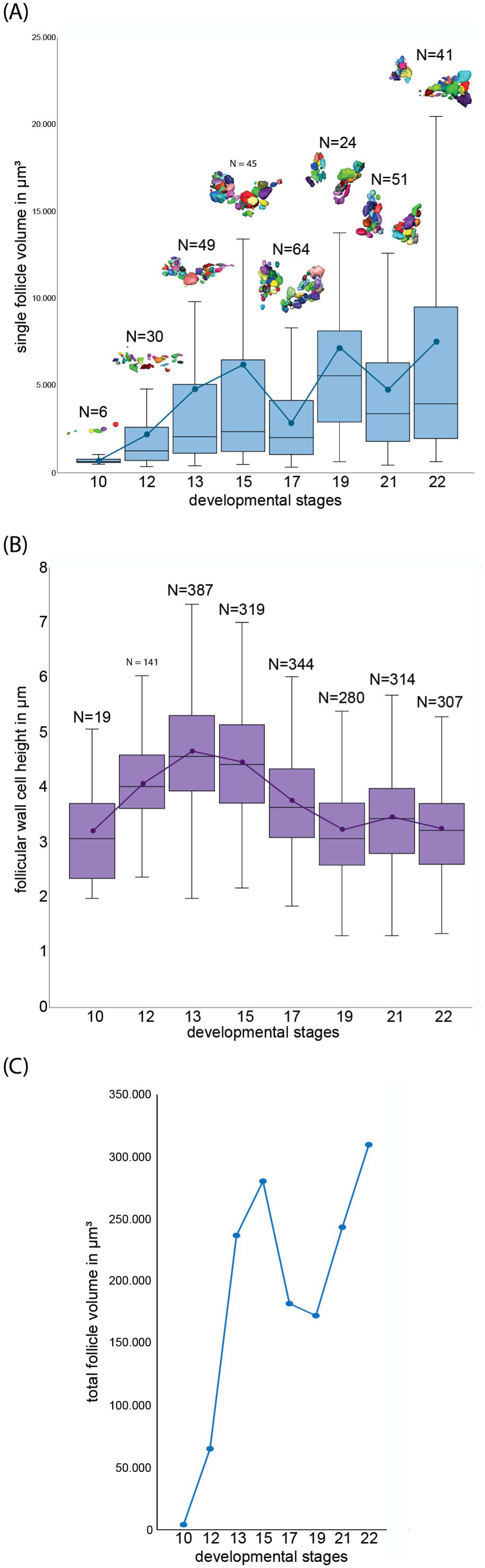
(A) Box plot of each individual follicle volume per developmental stage. Above each box is a 3-dimensional reconstruction of the follicles measured. Dark blue dots indicate the average follicle volume at each developmental stage and are connected via a line. (B) Diagram of the total follicle volume at each developmental stage. (C) Box plot of the follicle cell height at different developmental stages. Dark purple dots indicate the average follicle cell height at each developmental stage and are connected via a line.

**Table 1.** Summary of heterochronic shifts detected in the heterochrony plots.

| event/suite number | developmental events | timing in *X. laevis*<br>compared to *R. arenarum* | timing in *A. villiersi*<br>compared to *R. arenarum* | timing in *A. wahlbergii*<br>compared to *R. arenarum* | timing in *A. villiersi*<br>compared to *A. wahlbergii* |
| --- | --- | --- | --- | --- | --- |
| I | skin suite |  |  |  |  |
| 1 | Stratum compactum starts to develop | - | - | - | earlier |
| 2 | stratum compactum shows increased thickness | pre-displaced | - | post-displaced | earlier |
| 3 | intracellular yolk granules absent | - | - | - | - |
| 4 | dermal melanophores present | - | pre-displaced | pre-displaced | later |
| I.I | unicellular gland cell suite |  |  |  |  |
| 5 | unicellular mucus glands present | - | - | pre-displaced | later |
| 6 | unicellualr mucus glands absent | - | - | post-displaced | - |
| I.II | multicellular gland cell suite |  |  |  |  |
| 7 | Stratum spongiosum present | pre-displaced | - | - | - |
| 8 | multicellular gland anlagen | - | - | post-displaced | - |
| 9 | multicellular granular glands present | post-displaced | post-displaced | post-displaced | - |
| 10 | multicellular mucus glands present | - | - | - | - |
| I.III | epidermal cell layer suite |  |  |  |  |
| 11 | epidermis has up to two layers | - | - | - | - |
| 12 | epidermis has three or more layers | - | - | - | - |
| 13 | Stratum corneum present | pre-displaced | - | pre-displaced | - |
| II | hind limb suite |  |  |  |  |
| 14 | fist Anlage of the hind limb | - | pre-displaced* | pre-displaced | earlier |
| 15 | starting elongation of the hind limb buds | - | pre-displaced | pre-displaced | - |
| 16 | limb paddle present | - | - | - | - |
| 17 | toe 3-5 distinct | - | - | - | - |
| 18 | toe 1-5 present | - | - | - | - |
| III | forelimb suite |  |  |  |  |
| 19 | first Anlage of the forelimb buds | - | - | - | earlier |
| 20 | starting elongation of the forelimb buds | - | - | - | - |
| 21 | all 4 fingers differentiated | - | - | - | - |
| IV | tail suite |  |  |  |  |
| 22 | tail bud present | - | - | - | - |
| 23 | starting elongation of the tail bud | - | - | post-displaced | earlier |
| 24 | fin folds present | - | - | - | - |
| 25 | vascularized translucent fin folds | - | post-displaced | - | - |
| 26 | tail begins to regress | - | - | - | - |
| 27 | tail regressed to smal stump | - | - | - | - |
| 28 | tail resorbed | - | - | - | - |
| V | opercular fold suite |  |  |  |  |
| 29 | opercular fold first evident | - | - | - | - |
| 30 | opercular fold maximum extension | - | - | - | - |
| 31 | first perforation of th opercular fold | - | - | pre-displaced | later |
| 32 | both forelimbs are completely errupted | - | - | pre-displaced | later |
| VI | optic suite |  |  |  |  |
| 33 | optic bulges | - | - | - | earlier |
| 34 | pupil visible | - | - | - | - |
| 35 | dark eye pigment | - | - | - | - |
| 36 | coronoid fissure closed | - | - | - | - |
| 37 | eye lids present | - | - | - | - |
| VII | adult mouth suite |  |  |  |  |
| 38 | mouth slit before eye | - | pre-displaced | pre-displaced | - |
| 39 | mouth slit extends to the mid of the eye | - | - | pre-displaced | later |
| 40 | mouth slit behind the eye | - | - | - | - |
| VIII | thyroid gland suite |  |  |  |  |
| 41 | thyroid anlage present | - | - | - | - |
| 42 | follicle present | - | - | - | - |
| 43 | follicular wall cells present | - | - | - | - |
| 44 | follicular wall cells increse in thickness | - | - | - | - |
| 45 | follicular wall cells maximum thickness | - | - | - | later |
| 46 | collid present | - | - | - | - |
| IX | others |  |  |  |  |
| 47 | hatching | - | post-displaced | post-displaced | earlier |

**Table 2.** General developmental events that are used as criteria to define homologous ontogenetic phases compared in the heterochrony plots.

| #-Plot | ontogenetic phases | criteria (start of phase) | *Rhinella arenarum*<br>(Gosner-stage, 1960) | *Xenopus laevis*<br>(Nieuwkoop and Faber-stage, 1994) | *Arthroleptella villiersi*<br>(Schweiger-stage, 2017) | *Arthroleptis wahlbergii*<br>(Schweiger-stage, 2024) |
| --- | --- | --- | --- | --- | --- | --- |
| a | pre-pharyngula<br>(early embryonic) | fertilized egg | 1 | 1 | 0 | 1 |
| b | pharyngula and post-pharyngula<br>(late embryonic) | optic bulge; gill arches present;<br>tailbud elongated | 20 | 31 | 6 | 4 |
| c | "tadpole"<br>(pre- and prometamorphosis) | pupil visible, iris dark/black;<br>maximum extension of the opercular fold;<br>foot paddle-shaped | 25 | 45 | 8 | 7 |
| d | metamorphosis | begin of tail regression | 40 | 58 | 12 | 10,5 |
| e | postmetamorphosis | tail completely resorbed | 46 | 66 | 22 | 16 |

**Table 3.**
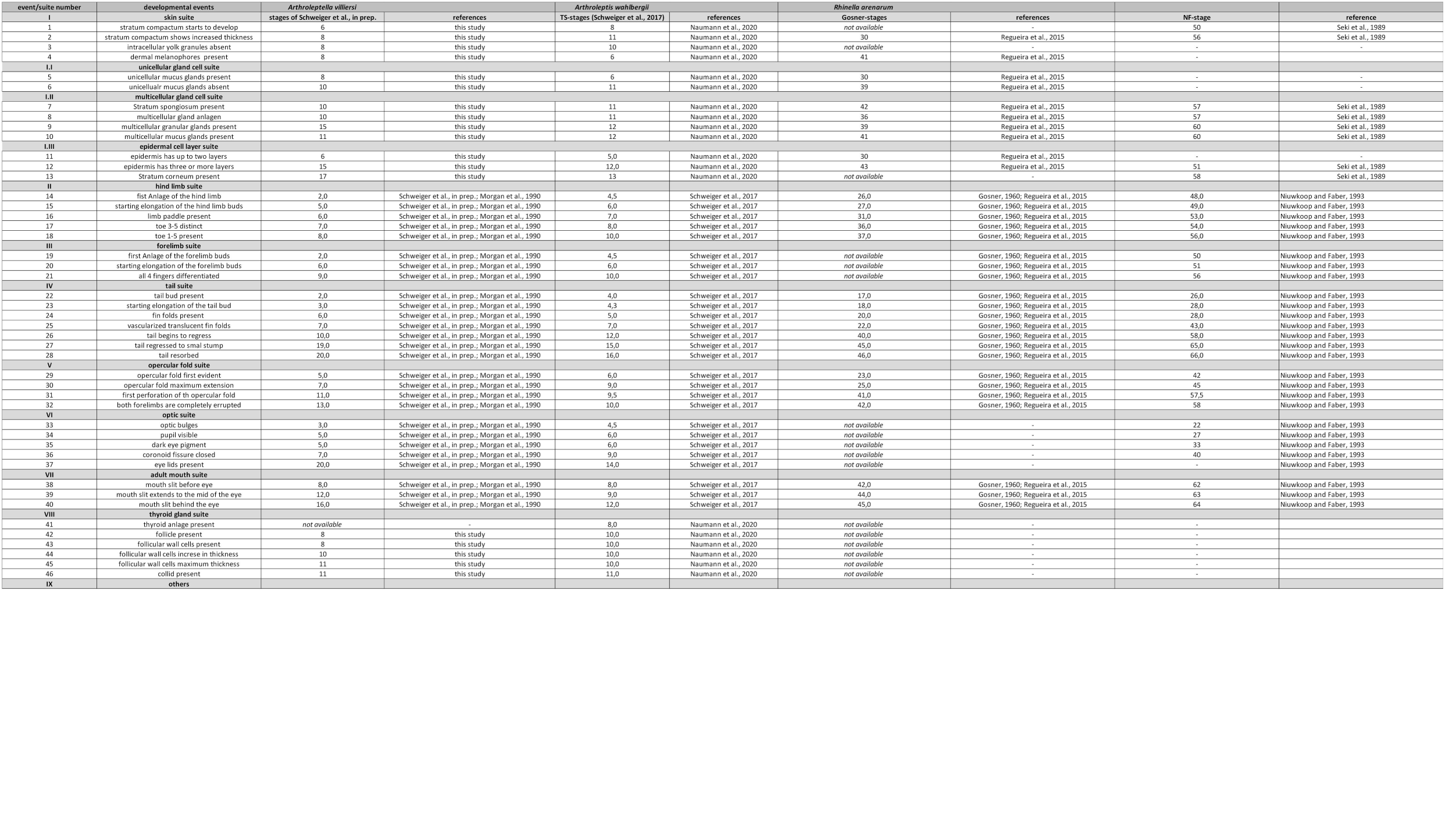
Developmental events investigated in the heterochrony plots.

## 3. Discussion

### 3.1 Melanophore patterning, skin development and metamorphic rearrangement correlate with thyroid gland maturation and are conserved among anurans with different life histories

Metamorphic changes in anurans, and amphibians in general, are initiated through the brain-pituitary-thyroid axis (BPT-axis), with TH as one important messenger to target tissues (Shi, 1999). In accordance with previous work on *Arthroleptella bicolor* Hewitt, 1926 (Brink, 1939), we could detect an early appearance of the thyroid gland (stage 10), an early and continuous presence of follicular colloid (from stage 12 on) and vascularization that is extensive at the beginning of thyroid differentiation (stage 12) but slowly decreases in later stages of *A. villiersi*. This pattern resembles the pattern reported for direct-developing *E. coqui* and *Arthroleptis* (Jennings and Hanken, 1998; Naumann et al., 2020). A correlation of BPT-axis maturation with overall skin development and metamorphic rearrangement has been shown in *A. lightfooti*, as well as in aquatic indirect- and terrestrial direct-developing taxa (Chammas et al., 2015; Fox, 1981, 1986; Goldberg et al., 2020; Morgan et al., 1989; Naumann et al., 2020; Regueira et al., 2016; Robinson and Heintzelman, 1987). The data from *A. villiersi* confirm the descriptions for *A. lightfooti* by Morgan et al. (1989) and add information on the exact timing of several developmental events (e.g., intracellular yolk granule resorption, dermal melanophore patterning, unicellular and multicellular gland complex differentiation; summarized in Figure 8). The anuran skin harbours several different types of chromatophores (Duellman and Trueb, 1994). However, in this study we focus on the pattern of melanophores only because they are the most accessible type using histology. In aquatic indirect-developing anurans, the tadpole-specific melanophore pattern develops first followed by the metamorphic changes of melanophore location and type, resulting in the adult pattern (Smith-Gill and Carver, 1981; Yasutomi, 1987). A comparable, albeit very condensed, melanophore rearrangement has been described for terrestrial direct-developing *Arthroleptis* (Naumann et al., 2020). In the terrestrial indirect-developing *A. villiersi*, melanophore patterning can be divided into two phases as observed in some other anurans (Naumann et al., 2020; Yasutomi, 1987). A first phase of increased pigmentation starts slightly before thyroid differentiation and is comparable to the establishment of the tadpole-specific coloration pattern. It is followed by a second phase of increased darkening of the skin that correlates with thyroid gland maturation and is comparable to the metamorphic rearrangement to the adult coloration pattern. We were not able to detect embryonic epidermal melanophores, or two melanophore populations with a distinctly different morphology, as described for other taxa (Naumann et al., 2020). The available data indicate that the two phases of melanophore patterning (tadpole-specific and postmetamorphic) are conserved among frogs with different reproductive modes. Furthermore, the second phase of melanophore patterning correlates with thyroid gland maturation and could hint at a TH dependency of the establishment of the adult colour pattern in *A. villiersi*. The development of a tadpole-like skin correlates with the timing of hatching (stage 8/9; stage 3 in Morgan et al., 1989). With the beginning of metamorphic rearrangement of the body skin, the proximal tail skin shows also signs of metamorphic rearrangements (e.g., developing multicellular glands and a stratum corneum). However, this is paralleled by tail degeneration in between stage 17 to 21. This indicates that the tail skin, similarly to the skin of the rest of the body, is competent to TH signalling, triggering at least the onset of metamorphic rearrangements. Postmetamorphic characters, like epidermal areas with three cell layers, a prominent stratum spongiosum, a stratum corneum, increased melanophore density and multicellular mucus and granular glands, develop from stage 13 to stage 19. In summary, skin development and metamorphic rearrangement appear conserved among anurans with different reproductive modes.

**Figure 8.**
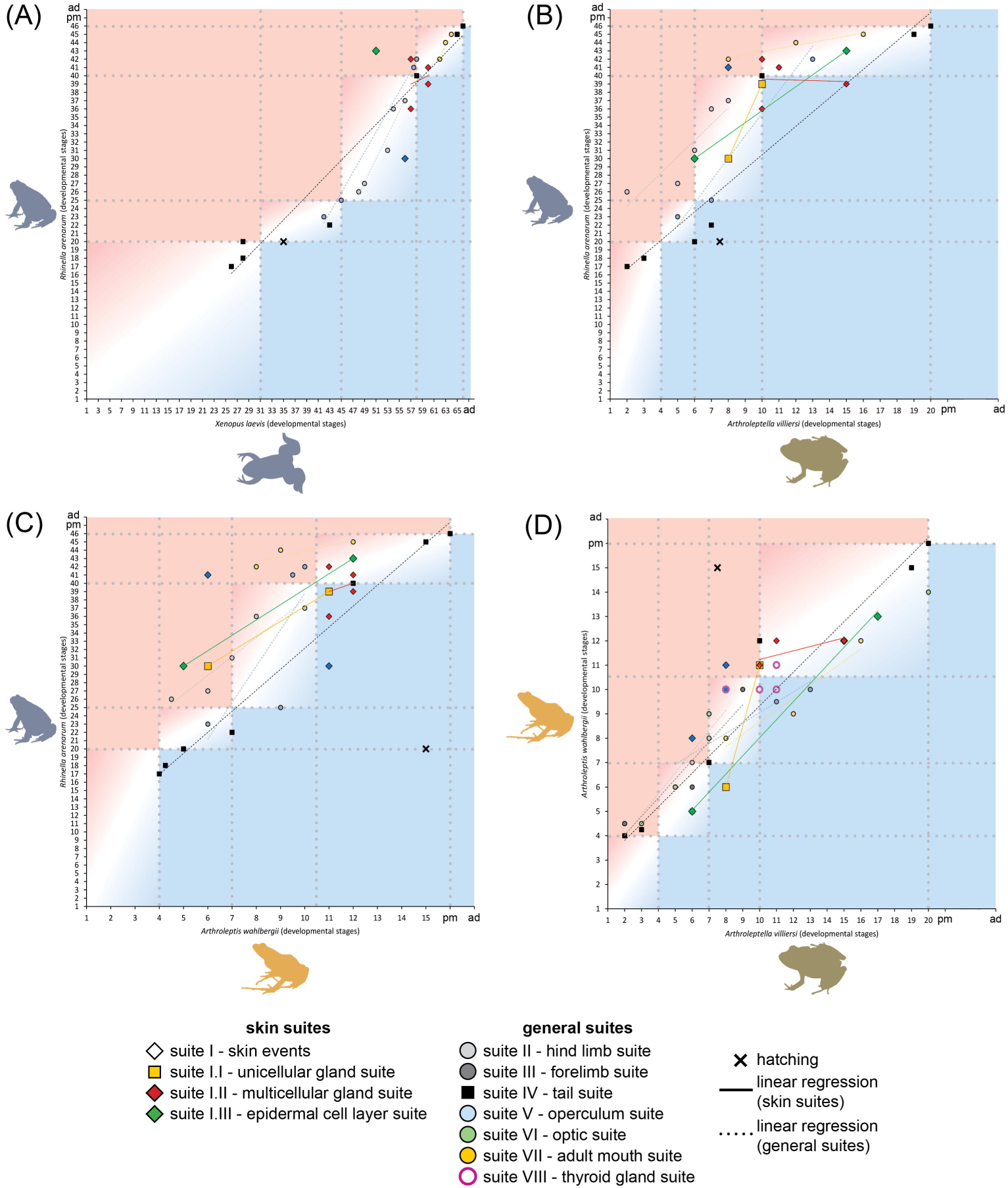
Heterochrony plots of anuran species with different reproductive modes. Light red background in the upper left area of a plot indicates pre-displacement/early development of an event, light blue background in the lower right area of a plot indicates post-displacement/later development of an event in the species on the x-axis. The white area indicates a comparable developmental timing.

**Figure 9.**
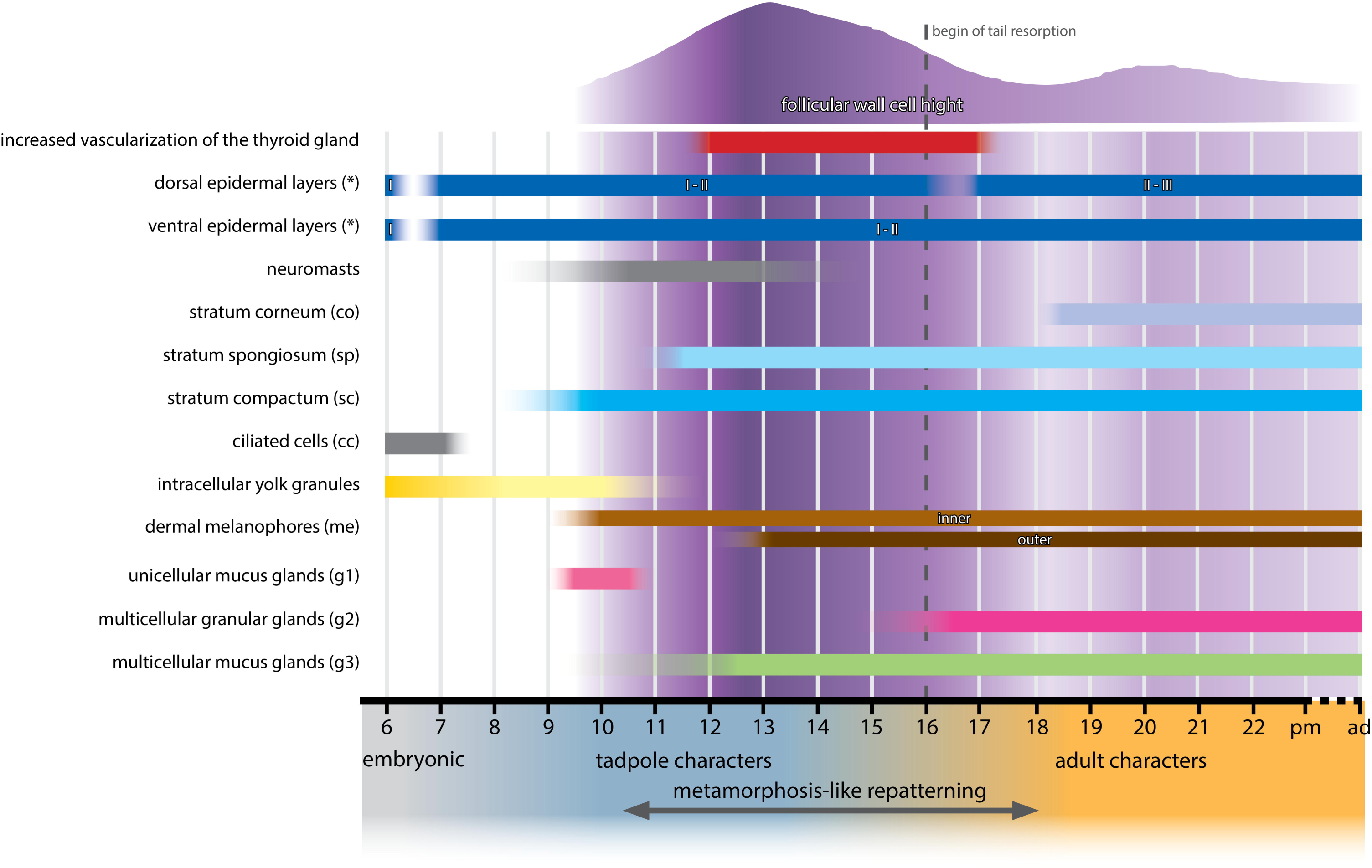
Schematic summary of the skin composition of *Arthroleptella villiersi* at different developmental stages. The histologically estimated thyroid gland activity based on follicle cell height measurements is indicated in different shades of purple. White to light purple indicates a lower activity while dark purple indicates higher activity.

### 3.2 Terrestrialization of development precedes neuromast loss in anurans

Previous studies have described the presence of neuromasts in *Arthroleptella* species but lacked detailed descriptions of their development and degeneration (de Villiers, 1929; Morgan et al., 1989). We confirm the presence of neuromasts in *A. villiersi* and describe their development in detail, starting with the first appearance of epidermal thickenings interpreted as placodes at stage 8/9, their differentiation until stage 12 (histologically similar to functioning neuromasts of aquatic indirect-developing species e.g. *Pseudis paradoxa* Linnaeus, 1758 (Quinzio and Fabrezi, 2014)), the beginning of their degeneration at stage 13, followed by their total absence at stage 15. The beginning of neuromast degeneration correlates with histological thyroid maturation. This pattern is similar to aquatic indirect-developing species (Wahnschaffe et al., 1987). It differs from the pattern in terrestrial direct-developing *Arthroleptis* and *Eleutherodactylus* in which neuromasts never develop (Naumann et al., 2020; Schlosser et al., 1999; Townsend and Stewart, 1985). We propose two hypotheses to explain the conserved developmental pattern of neuromast in *A. villiersi*. (1) The presence of neuromasts is advantageous for hatchlings to sense movements of the surrounding fluid of the jelly nest. However, there is only a very short ontogenetic period where possibly functional neuromasts are present. (2) The few neuromasts are remnants, possibly even non-functional, representing an evolutionary intermediate state on the way to the complete loss of this sensory organs. The second hypothesis can be tested by investigating the presence of neuromast-associated components of the central and peripheral nervous system in *A. villiersi*. The observed pattern of the short phase of neuromast presence in *A. villiersi* and their complete absence in terrestrial direct-developing taxa support previous interpretations that neuromasts, or the lateral line in general, represents a distinct developmental module (Schlosser et al., 1999).

### 3.3 Heterochronic shifts of skin development in anurans with different life histories indicate a modular organization of the anuran skin

By comparing the sequences of a set of general metamorphic events (hindlimb development, degeneration of the pronephros, skin metamorphosis, forelimb emergence, tail development and degeneration) with the timing of exogenous feeding, hatching and BPT-axis maturation between *A. lightfooti, Xenopus* and *Eleutherodactylus*, Morgan et al. (1998) concluded that the developmental pattern of *A. lightfooti* is temporally intermediate between the direct-developing *Eleutherodactylus*, and the aquatic indirect-developing *Xenopus*. However, no skin developmental data for *Eleutherodactylus* or any other direct-developing species where available for this previous study. We constructed heterochrony plots containing 13 homologous skin developmental events and 34 general developmental events grouped into nine suites (Roman numerals) to test the predictions of Morgan and colleagues. The heterochrony plots in general allow the view of the developmental pattern of *Arthroleptella* as an intermediate pattern between aquatic indirect and terrestrial direct development as stated by Morgan et al (1998). This is demonstrated by (1) dermal melanophores appearing earlier in terrestrial reproducing species investigated and first in *Arthroleptis*, (2) a similar development of unicellular mucus glands in indirect developing species investigated vs. an early appearance and long persistence in the direct developing *Arthroleptis,* (3) a post-displacement of multicellular granular gland differentiation in terrestrial reproducing species investigated, (4) a pre-displaced development of the stratum corneum in *Arthroleptis* compared to *Arthroleptella*.

Previous studies on heterochronic shifts during the early ontogeny (early tailbud-stage until the “complete tadpole”) in various anuran groups proposed formalized heterochrony analyses as a promising road to identify developmental modules (Grosso et al., 2019; Grosso et al., 2020; Grosso et al., 2017; Thibaudeau and Altig, 1988; Vera-Candioti et al., 2016). Following this program by preparing heterochrony plots developed by Schlosser and Roth (Schlosser, 2001; Schlosser and Roth, 1997), the present study indicates that the anuran skin is constructed of different developmental modules. If these modules tend to be modified and repeatedly co-dissociate from other structures (i.e. exhibit coordinated heterochronic shifts), such as the developing tail limbs, they should also act as evolutionary units (Schlosser et al., 1999). A developmental module showing strong evidence for representing an evolutionary unit is the neuromast/lateral line module. Neuromasts, normally present during the tadpole phase and degenerating during metamorphosis, can be lost in direct-developing taxa (Naumann et al., 2020; Schlosser et al., 1999; Townsend and Stewart, 1985) but are usually maintained during the adult phase in aquatic anurans (Fabrezi and Quinzio, 2008; Nieuwkoop and Faber, 1956), and their organization and number within distinct lateral lines can be altered (Lannoo, 1987) without changing the overall organization of the skin. Following a similar reasoning it should be possible to identify additional developmental modules within the anuran skin that might represent evolutionary units. Following a similar reasoning it should be possible to identify additional developmental modules within the anuran skin that might represent evolutionary units. The heterochronic pattern of some skin developmental events indicate the presence of dermal melanophore, unicellular and at least two multicellular gland developmental modules (see Table 1). However, if these modules coevolve as one evolutionary unit (e.g., a general skin evolutionary unit is hampered by the scarcity of comparative skin developmental data and a lack of investigations of mechanistic data on anuran skin development.

## 4 Experimental procedures

### 4.1 Specimens

Embryos, hatchlings and adults of *A. villiersi* were collected in different locations of the Western Cape, South Africa. Clutches were transferred to petri dishes and maintained under field conditions at ambient temperatures, and embryos sampled at regular intervals (every 1-2 days). Specimens were euthanized using tricaine methanesulfonate (MS222; Fluka), fixed in 4% buffered formalin, and subsequently stored in 70% ethanol. Conspecificity of the egg clutches was confirmed by raising individual eggs to hatching and through the guarding male (du Preez and Carruthers, 2017). For each developmental stage examined in this study, one individual was examined externally to describe the pigmentation pattern and another individual was sectioned to describe histological changes of the skin. For further details on specimen collection and staging see Schweiger et al. (2025).

### 4.2 Photographs and Image processing

Photographs of whole specimens were taken using a Zeiss Discovery V12 stereomicroscope equipped with a Zeiss AxioCam. Brightness and contrast were adjusted, and background noise was removed using Adobe Photoshop CS6.

### 4.3 Histological sectioning and staining

Embryos were prepared for sectioning as described for *Arthroleptis* (Naumann et al., 2020). Specimens at developmental stages 4 and 6/7 were embedded in Technovit (Kulzer R0010022), sectioned at 3 µm and stained with a mixture of basophilic Methylene blue and acidic Fuchsin (Böck, 1989). Embryos from stage 8 on were decalcified for up to three days (Osteomoll, Merck), embedded in Paraplast (Roth, X881.1), sectioned at 7 µm and stained either with Heidenhain’s Azan or Goldner’s Trichrom (Böck, 1989). All sections were prepared with a Microm HM 360 (Zeiss). Photographs were taken using a Leica DM750.

### 4.4 Histological measurements

Measurements and 3-dimensional reconstructions of single follicle cell volumes were obtained from digitized and aligned serial sections using Amira 5.4.2 (FEI Visualization, Science Group, Bordeaux, France). Follicle cell heights were measured from digitized sections using ImageJ (Schneider et al., 2012). Diagrams were prepared using Microsoft Excel 2016.

### 4.5 Heterochrony plots

In heterochrony plots, the timing of developmental events in one taxon (x-axis) is plotted against the timing of homologous events in another taxon (y-axis) (Schlosser, 2001; Schlosser & Roth, 1997). In this study, 47 developmental events (13 developmental events of the skin and 35 general events) were conceptualized. Causally linked events were grouped into nine developmental suites indicated by Roman numerals. Events were excluded from the plot between two taxa when data on the developmental timing were only available for one taxon. Linear regression lines are given for the suites to estimate their developmental rates. To facilitate a more comprehensive comparison, five ontogenetic phases were defined based on very general developmental events and indicated by dotted lines in each plot. Phase a (“pre-pharyngula” phase): fertilization until the last stage before the emergence of optic and gill arch bulges. Phase b (“pharyngula and post-pharyngula” phase): appearance of optic and gill arch bulges as well as the elongation of the tail bud until the last stage before the visibility of the pupil, the darkening of the iris as well as the maximum extension of the operculum (additionally the presence of forelimb buds beneath the operculum and a paddle-shaped foot in direct-developing species). Phase c (“tadpole” phase): first appearance of the of the pupil, the darkening of the iris as well as the maximum extension of the operculum (additionally the presence of forelimb buds beneath the operculum and a paddle-shaped foot in direct-developing species) until the last stage before first signs of tail regression. Phase d (“metamorphosis” phase): onset of tail regression until one stage before the completion of metamorphosis defined by the complete disappearance of a tail stub. Phase e (“postmetamorphosis” phase): complete absence of a tail stub onwards (see Table 2 for specific stages).

## Supporting information

Supplementary Material 1. Histological measurements of the thyroid gland of different developmental stages of Arthroleptella villiersi.

## Acknowledgements

We would like to thank Katja Felbel for valuable help in the laboratory. Collection of material was facilitated by Alan Channing and Andrew Turner, whose advice, help in the field, and hospitality are gratefully acknowledged. Permits to collect were issued by CapeNature (permits No. 0035-AAA004-01041, 0056-AAA041-00073). Fieldwork was funded through a German Science Foundation (DFG) grant to HM (MU 2914/2-1) to HM, which is gratefully acknowledged.

**Supplementary Material 1.** Histological measurements of the thyroid gland of different developmental stages of *Arthroleptella villiersi*.

