## Supplementary figures and images for "Skin development and metamorphic re-arrangement in *Arthroleptella villiersi* Hewitt, 1935 reveals distinct heterochronic patterns in anurans with different life histories"

### Supplementary Material 1. Histological measurements of the thyroid gland of different developmental stages of Arthroleptella villiersi.

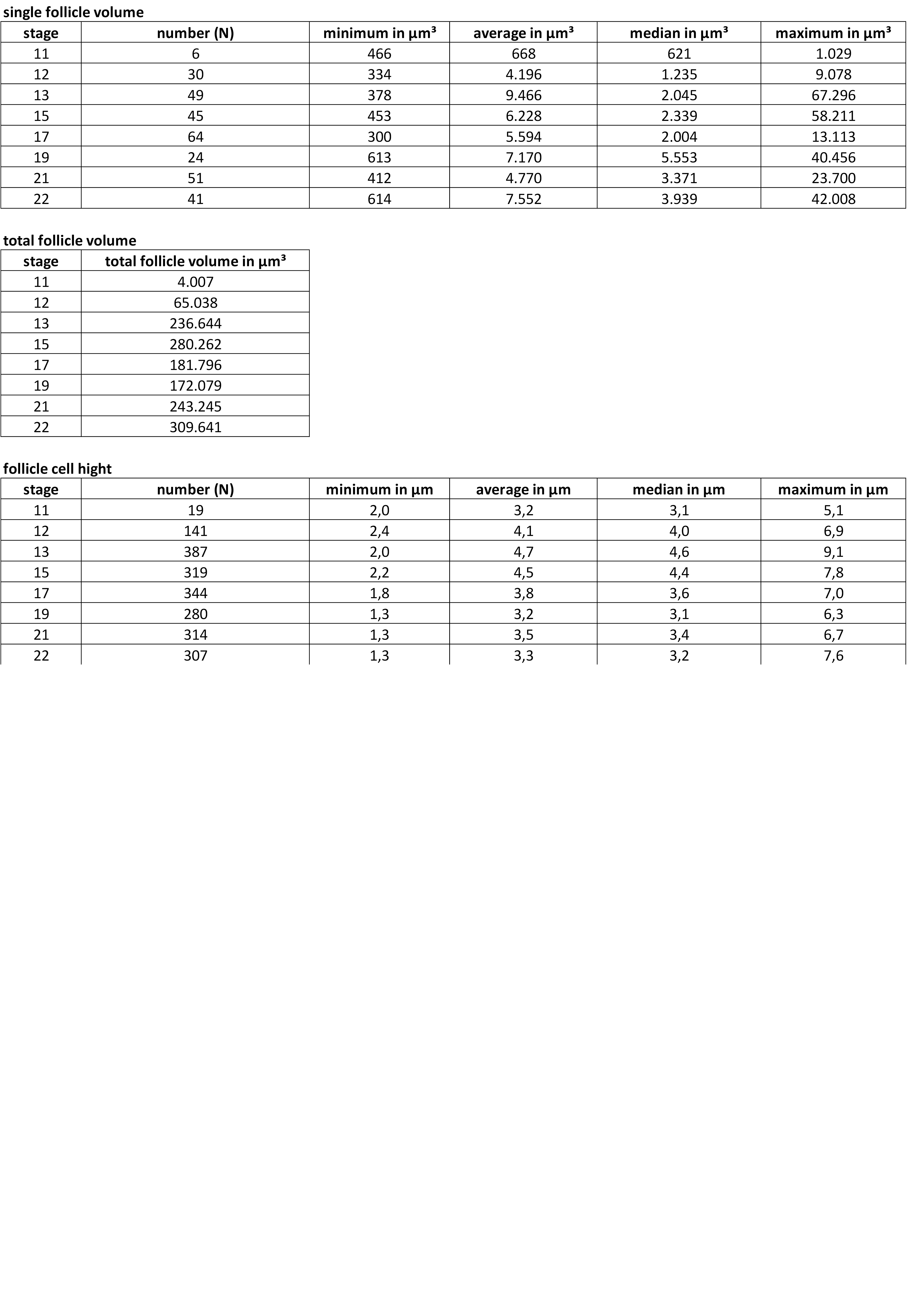
